# A Critical, Nonlinear Threshold Dictates Bacterial Invasion and Initial Kinetics During Influenza

**DOI:** 10.1101/052175

**Authors:** Amber M. Smith, Amanda P. Smith

## Abstract

Secondary bacterial infections increase morbidity and mortality of influenza A virus (IAV) infections. Bacteria are able to invade due to virus-induced depletion of alveolar macrophages (AMs), but this is not the only contributing factor. By analyzing a kinetic model, we uncovered a nonlinear initial dose threshold that is dependent on the amount of virus-induced AM depletion. The threshold separates the growth and clearance phenotypes such that bacteria decline for dose-AM depletion combinations below the threshold, stay constant near the threshold, and increase above the threshold. In addition, the distance from the threshold correlates to the growth rate. Because AM depletion changes throughout an IAV infection, the dose requirement for bacterial invasion also changes accordingly. Using the threshold, we found that the dose requirement drops dramatically during the first 7d of IAV infection. We then validated these analytical predictions by infecting mice with doses below or above the predicted threshold over the course of IAV infection. These results identify the nonlinear way in which two independent factors work together to support successful post-influenza bacterial invasion. They provide insight into coinfection timing, the heterogeneity in outcome, the probability of acquiring a coinfection, and the use of new therapeutic strategies to combat viral-bacterial coinfections.

## Introduction

Influenza A virus (IAV) poses a considerable threat to public health, resulting in 15-65 million infections and >200,000 hospitalizations each year during seasonal epidemics in the U.S.^1, 2^. Morbidity and mortality increase when a pandemic strain emerges and/or when IAV infection is complicated by a bacterial pathogen like *Streptococcus pneumoniae* (pneumococcus), which has accounted for 40-95% of influenza-related mortality in past pandemics^3-6^. As the respiratory tract environment deteriorates during influenza, the physiological barriers and immune mechanisms that normally clear pathogens become compromised and bacteria are able to invade and grow rapidly. Several factors, including viral and bacterial strain, inoculum size, and bacterial infection timing, are thought to contribute to influenza-bacterial coinfection kinetics, pathogenicity, and the likelihood of severe pneumonia developing (reviewed in^7-13^). Understanding how each factor influences the virulence and interaction between influenza viruses and bacterial pathogens and how each is interrelated is pivotal to finding effective preventative and therapeutic strategies.

Although well-characterized animal models have allowed for the study of various factors that affect bacterial acquisition and pathogenicity after influenza (reviewed in^11^), the extraordinary complexity of host-pathogen and pathogen-pathogen interplay complicates investigating every possible interaction simultaneously. Quantitative analyses have made it possible to simultaneously assess the contributions of different components and identify critical mechanisms driving influenza-bacterial coinfection kinetics. We recently combined a mathematical model and data from animal studies to establish dynamical host-pathogen feedbacks, quantify the contribution of various hypothesized mechanisms (e.g., virus enhanced bacterial attachment^14, 15^ and alveolar macrophage (AM) inhibition^16^), and develop new hypotheses (i.e., bacteria enhanced virus production) about the relationship between influenza and pneumococcus^17^. Our mathematical model revealed that the rapid increase in bacterial loads, a hallmark of influenza-pneumococcal coinfection, is initiated by the virus removing the protective effect of alveolar macrophages (AMs) with 85-90% efficiency by 7d post-influenza infection (pii) and that bacterial clearance could be achieved with improved AM response. This was in correlation to one experimental study suggesting that the phagocytic ability of these cells is inhibited^16^. It was initially unclear from either study if the effect accumulates over time and if it comprises several underlying mechanisms. However, a more recent experimental study followed up these works by using an advanced gating strategy to better define the AM population throughout the course of an IAV infection and found that these cells are depleted^18^, rather than or in addition to their functional inhibition, by IAV and that the level of depletion corresponds to the amount of bacterial outgrowth^18^. While our mathematical model does not distinguish between these mechanisms, the AM data indicated that the maximum amount of depletion occurred 7d pii and matched our parameter estimate of 85-90%^18^. Further, our model did include a handling time effect on the rate of bacterial phagocytosis by AMs, which had only a minor role. This further supports AM depletion as the dominant mechanism driving bacterial establishment, with functional inhibition as a possible secondary mechanism, and the accuracy of our model. Remarkably, this also corresponds to the time when bacterial coinfections are the most lethal^19^. The underlying mechanism resulting in the loss of AMs during influenza virus infection is currently unknown.

Another important feature of influenza-pneumococcal coinfection biology is that bacteria grow rapidly for initial doses that would be rapidly cleared in the absence of virus^17, 19^. In both naive and influenza-infected hosts, the trajectory of bacterial titers is dependent on the inoculating dose^16, 17, 20-22^. Further, in the context of the coinfection, a distinct dichotomous pattern emerged with a low dose (10^2^ CFU) compared to a higher dose (103 CFU) such that some individuals had high bacterial titers, an indication of severe pneumonia, while others had low bacterial titers and, presumably, a more mild infection^17^. It is currently unclear what factors contribute to the differential dynamics, although we hypothesized that this may be due to heterogeneity in the AM population because reducing the depletion parameter in our mathematical model could result in lower titers^17^ and because we previously related the dose dependent effect to the AM population in a naive host^20^. However, the exact connection between dose and AM depletion during influenza-pneumococcal coinfection is unclear. Understanding this behavior in more detail will help elucidate what conditions lead to secondary pneumococcal infections after influenza and why only a proportion of IAV infections lead to severe bacterial pneumonia^23-26^.

Here, we analyzed our coinfection model in more detail to quantify how the trajectory of bacterial growth changes with the level of AM depletion. In doing so, we uncovered a nonlinear initial dose threshold that is dependent on the number of AMs and found the critical number of AMs required to support a clearance phenotype. We then used this threshold together with data on the AM population^18^ to predict how the dose requirement declines over the course of an IAV infection. To validate the time-dependent threshold predictions, we examine bacterial growth/clearance rates in groups of BALB/cJ mice infected with influenza A virus A/Puerto Rico/8/34 (H1N1) (PR8) then 1, 3, 5, 7, 9, or 11 days later with pneumococcal strain D39 at a dose larger or smaller than the predicted threshold. The resulting data agree with our theoretical predictions and, in doing so, define an important, nonlinear relationship between bacterial dose and AM depletion that can be used as a predictive tool. Taken together, our data gives new insight into the timing of coinfections, potential therapeutic strategies, and why bacterial coinfections occur more often during influenza pandemics compared to seasonal epidemics.

## Results

### Coinfection Kinetics Depend on AM Depletion

To examine how pneumococcal kinetics during IAV infection change with varying degrees of AM depletion, we simulated the coinfection model (Equations (2)-(6)) with values of AM depletion (*ϕ*) ranging between 0% and 100% (0 ≤ *ϕ* ≤ 1). The resulting dynamics (Figure 1A) illustrate that distinct bacterial outcomes (i.e., maximum growth or clearance) exist depending on the degree that the IAV infection reduces the AM population. Bacterial resolution is predicted to occur with minor AM depletion (small *ϕ*), but the length of time for bacterial loads to completely clear (log10 *P*(*t*) < 0, where *P*(*t*) denotes the solution to Equation (6)) increases rapidly as depletion accumulates (increasing *ϕ*) and saturates once these cells have declined by ~80% (Figure 1B).

**Figure 1.**
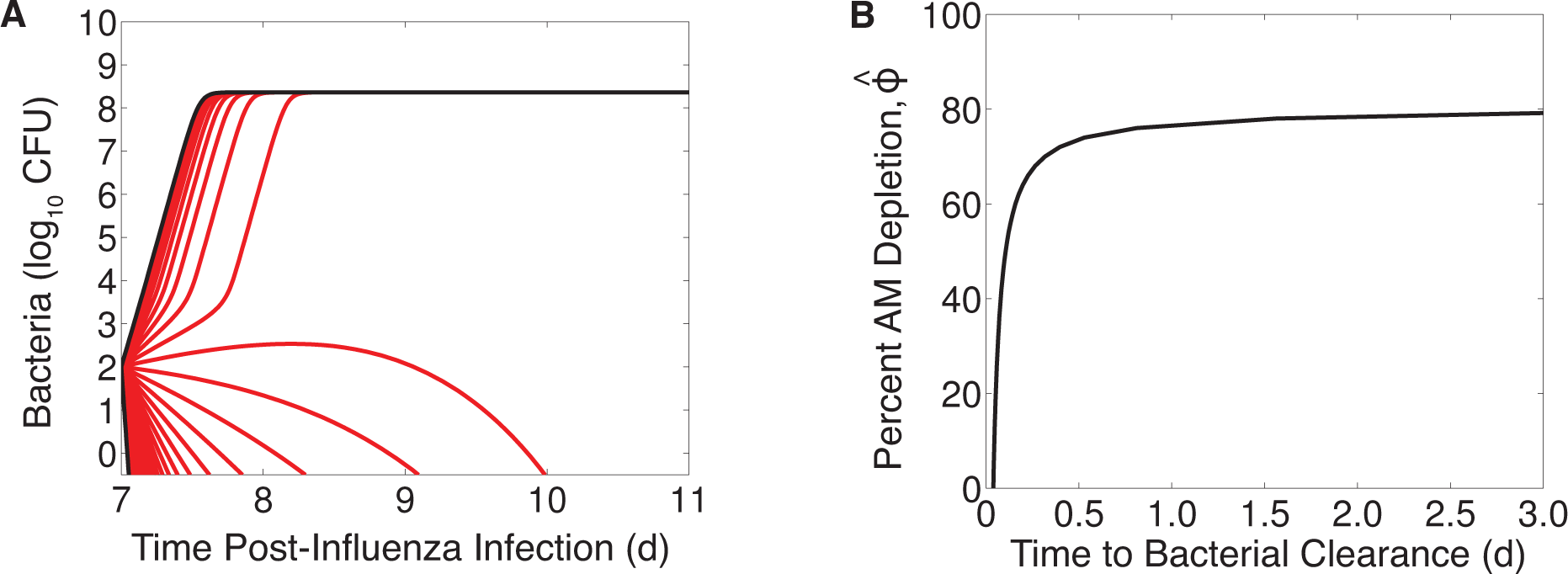
Simulation of the coinfection model for various levels of AM depletion. (A) Simulation of the coinfection model (Equations (2)-(6)) for infection using the parameter values in Table S1 that were optimized for an infection with 10^2^ TCID50 PR8 followed 7d later by 103 CFU D39 and values of AM depletion (*ϕ*) ranging from no impairment (*ϕ* = 0), which results in immediate clearance, to 100% impairment (*ϕ* = 1), which results in immediate growth to the maximum carrying capacity. (B) The number of days until complete bacterial clearance (log10 *P*(*t*) < 0, where *P*(*t*) is the solution to Equation (6)) occurs for AM depletion between 0% and 100%.

### Initial Dose Threshold

Because different levels of AM depletion result in different outcomes, we analyzed our coinfection model using mathematical steady-state and bifurcation analyses (details in the Supplementary Information). This analysis verified the two potential outcomes (stable steady states) as clearance (*P* = 0 CFU) and sustained growth to maximum carrying capacity (*P* = *K_P_* = 2.3 × 10^8^ CFU) (Figure 2A). An intermediate, unstable state given by Equations (7)-(8) is dependent on the extent of AM depletion and governs which of these outcomes manifests. This state predicts an initial dose threshold (Figure 2A) such that bacteria will exhibit a growth phenotype for doses above the threshold and a clearance phenotype for doses below the threshold for a given amount of AM depletion. Further, the rate at which bacteria grow or clear will increase as the distance from the threshold increases. As AMs become more depleted, the dose needed to elicit an infection drops rapidly in a nonlinear manner. Once AMs are reduced by ~80%, as indicated by the near zero value of the threshold, any dose will support bacterial growth. This critical level of depletion (*ϕ̂_crit_*) can be analytically calculated (see the Supplementary Information) in terms of the model parameters (Table S1) as

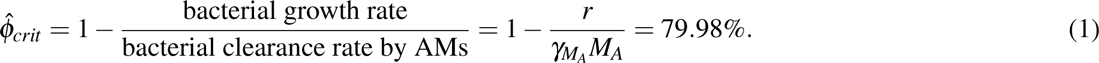

**Figure 2.**
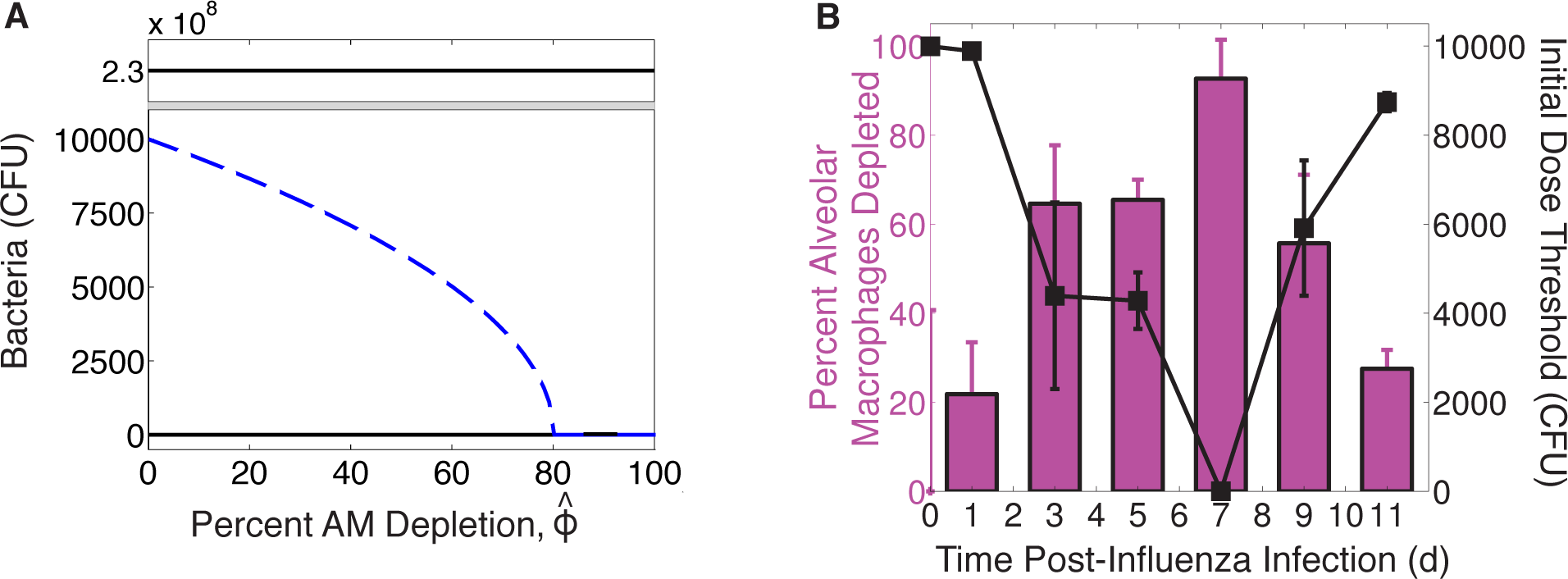
Initial dose threshold dependence on AM depletion. (A) Bacterial outcomes (steady states) for the coinfection model for different values of AM depletion (*ϕ̂*). Two possible outcomes exist, clearance (0 CFU) or growth to maximum carrying capacity (*KP* = 2.3 × 10^8^ CFU), and are indicated by solid black lines. The intermediate state (dashed blue line, defined by Equations (7)-(8)) is the initial dose threshold that dictates if bacterial loads will decline (doses below the threshold) or if they will increase (doses above the threshold). (B) The predicted threshold (black line, calculation in Table 1) over the course of an IAV infection given the amount of AM depletion^18^, Copyright 2013. The American Association of Immunologists, Inc. (magenta bars; see Methods).

### Threshold Dynamics During IAV Infection

Because AM depletion is dynamic during IAV infection^18^ and the initial dose threshold is dependent on AM depletion, the initial dose threshold will change throughout the course of an IAV infection. To determine how the threshold changes with time, we used *in vivo* data on the number of AMs lost at 1, 3, 5, 9, and 11d pii (see Methods) (Figure 2B) as an approximation for the parameter *ϕ* in Equation (6) because our estimated parameter value^17^ matched the empirical value^18^ for a coinfection at 7d pii. We used the mean and the standard deviation of the AM data to obtain an estimated confidence interval for *ϕ* for each time point (Table 1). We then used these values together with the model solution for virus (*V*(*t*)) at each time point to calculate the overall effect of depletion, defined by *ϕ̂* = *ϕV*/(*K_PV_* + *V*) in Equation (6), and the corresponding initial dose threshold value at each coinfection time (Table 1). The resulting threshold drops rapidly between 1d and 3d, reducing the dose necessary for a secondary infection to establish by over 50% (Figure 2B). The threshold decreases by another 50% by 7d pii before increasing to near baseline level at 11d pii.

**Table 1.** Calculation of the estimated initial dose threshold. The threshold (Equations (7)-(8)) is calculated using the overall AM depletion estimate (*ϕ̂* = *ϕV*(*t*)/(*K_PV_* + *V*(*t*))), where *ϕ* is estimated by the percent AM depletion^18^ (Copyright 2013. The American Association of Immunologists, Inc.; Figure 2B) and *V*(*t*) is the coinfection model solution for virus.

| Coinfection Day | Percent AM Depletion Estimate of $\phi$ | Virus, $\log_{10}$ V(t) Model Solution<sup>17</sup> ($\log_{10}$ TCID<sub>50</sub>) | Percent Overall AM Depletion $\hat{\phi} = \phi \frac{V(t)}{K_{PV} + V(t)}$ | Initial Dose Threshold Equations (7)-(8), $\times 10^3$ (CFU) |
| --- | --- | --- | --- | --- |
| 1 | 21.8 [10.2, 33.5] | 2.2 | 1.9 [0.02, 19.5] | 9.89 [9.83, 9.94] |
| 3 | 64.6 [51.4, 77.8] | 6.7 | 64.6 [50.9, 77.8] | 4.39 [2.29, 6.49] |
| 5 | 65.4 [60.9, 70.0] | 6.1 | 65.3 [58.7, 70.0] | 4.28 [3.65, 4.91] |
| 7 | 92.7 [83.9, 100] | 5.3 | 91.8 [43.2, 100] | $2.2 \times 10^{-5}$ [$4.3 \times 10^{-6}$, $5.2 \times 10^{-5}$] |
| 9 | 55.7 [40.3, 71.1] | 4.5 | 52.9 [0.67, 71.1] | 5.91 [4.39, 7.43] |
| 11 | 27.5 [23.3, 31.7] | 3.7 | 20.9 [0.05, 31.7] | 8.74 [8.54, 8.94] |

### Bacterial Growth for Doses Below or Above the Threshold

The initial dose threshold indicates that bacteria will clear for doses lower than the threshold and grow for doses higher than the threshold. To test the predicted dynamic threshold, we infected groups of mice first with 50 TCID_50_ PR8 then D39 at 1, 3, 5, 7, 9, or 11d pii at a bacterial dose below or above the predicted threshold value (Table 2). For bacterial infection 7d pii, a dose below the threshold was not examined due to the predicted value being less than 1 CFU. We then quantified bacterial growth at 4h and 24h post-bacterial infection (pbi).

**Table 2.** Bacterial doses used to test the predicted initial dose threshold.

| Coinfection Day | Threshold $\times 10^3$ (CFU) | Dose Below Threshold $\times 10^3$ (CFU) | Dose Above Threshold $\times 10^3$ (CFU) |
| --- | --- | --- | --- |
| 1 | 9.89 | 3.50 | 10.7 |
| 3 | 4.39 | 2.75 | 9.40 |
| 5 | 4.28 | 2.70 | 5.90 |
| 7 | $2.2 \times 10^{-5}$ | - | 0.50 |
| 9 | 5.91 | 3.90 | 8.90 |
| 11 | 8.74 | 3.65 | 10.0 |

When inoculating with a dose below the predicted threshold, we found that bacterial loads at 4h pbi were significantly lower (p<0.05) than the inoculum in all mice for all coinfection times tested (Figure 3A). This initial clearance supports our predicted threshold, which suggests that doses below the threshold will result in bacterial clearance. For a coinfection at 1d pii or 3d pii and for the doses used, bacteria were undetectable in 2 out of 5 mice. To investigate whether clearance could also be attained with a dose closer to the threshold value, we inoculated a group of influenza-infected mice with 7.55 × 10^3^ CFU (compared to 3.55 × 10^3^ CFU) D39 at 1d pii. At this dose, bacterial loads decreased in all mice (p<0.05) but none resolved the infection within 4h pbi. Conversely, we also examined a dose lower than the low dose in Table 2 (1 × 10^3^ CFU compared to 2.7 × 10^3^ CFU) for bacterial infection 5d pii and found that the rate of clearance improved and that 1 out of 5 mice resolved the infection (p<0.05).

**Figure 3.**
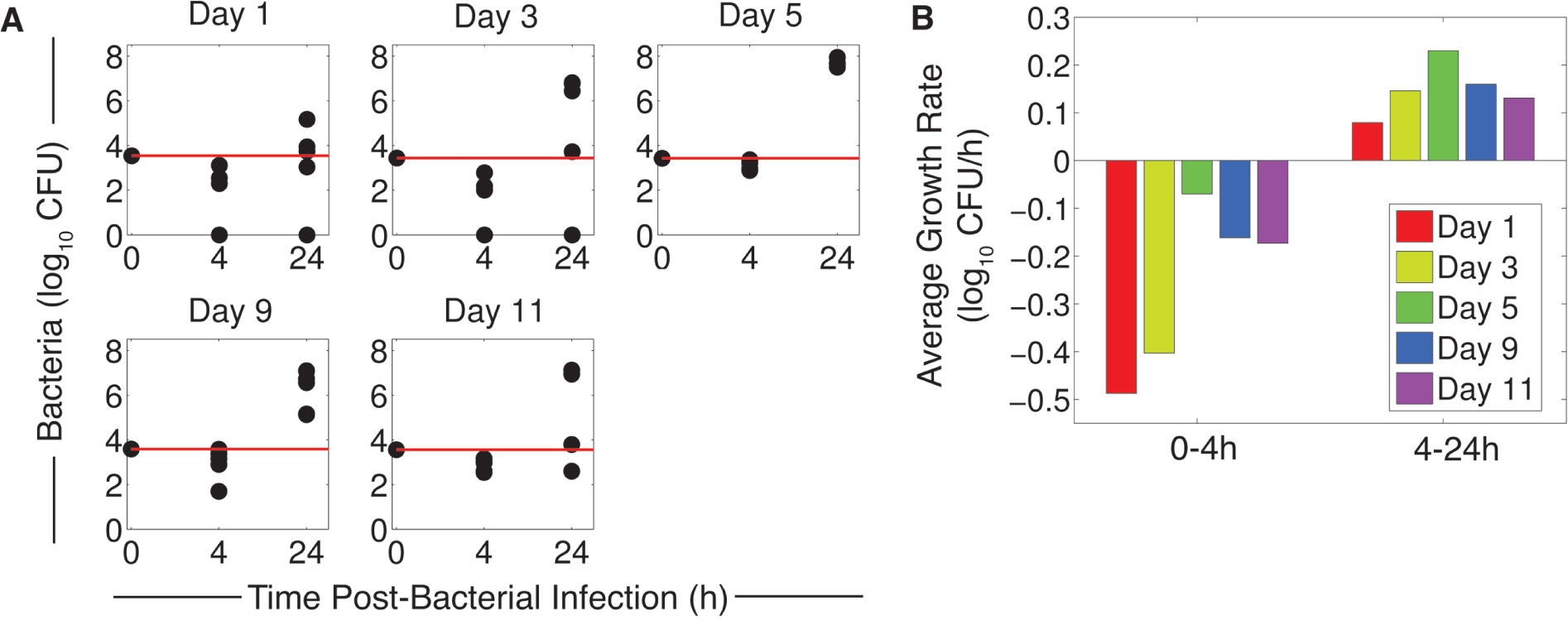
Bacterial growth for doses below the threshold. (A) Bacterial loads (log10 CFU) at 4h and 24h for PR8 infection followed by D39 at the indicated coinfection time (1, 3, 5, 9, or 11d pii) at doses below the threshold (Table 2). The red line indicates the inoculum size and each dot represents an individual mouse (5 mice/group). (B) Average log10 rate of bacterial growth from 0h to 4h and from 4h to 24h for coinfection at 1, 3, 5, 9 or 11d post-influenza at the indicated coinfection time.

We next examined bacterial titers at 24h pbi in a separate group of mice using the low doses in Table 2 to determine if titers would continue to decline. For coinfection at 1d and 3d pii, where 40% had resolved the infection within 4h, the same proportion of individuals had undetectable titers at 24h pbi. If bacteria were not controlled within this time frame, then the initial clearance mechanisms were overcome and bacterial loads surpassed those at 4h and, in some cases, the inoculum (Figure 3A). The average rate of growth between 4h and 24h inversely correlates to the average clearance rate between 0h and 4h such that slower clearance supports faster growth (Figure 3B) and less titer heterogeneity (Figure 3A).

When mice were given a dose slightly above the threshold, bacterial loads at 4h pbi remained relatively constant (p*>*0.05) and showed dichotomous behavior for all coinfection timings with 50-60% of individuals partially cleared the inoculum and the remaining 40-50% showed rapid growth (Figure 4A). To find the dose where 100% of infections result in immediate growth and to show that growth increases with the distance from the threshold, we infected groups of mice at 5d pii with doses incrementally higher than in Table 2 (7.0, 8.5, or 10.5×10^3^ CFU) (Figure 4B-C). At 7 × 10^3^ CFU, some individuals were still able to partially clear the inoculum (p>0.05). However, at 8.5 × 10^3^ CFU and 10.5 × 10^3^ CFU, bacteria readily grew in all individuals to levels higher than the inoculum (p<0.005). In addition, the strength of this growth increased and the titer heterogeneity decreased as the dose increased (Figure 4C-D).

**Figure 4.**
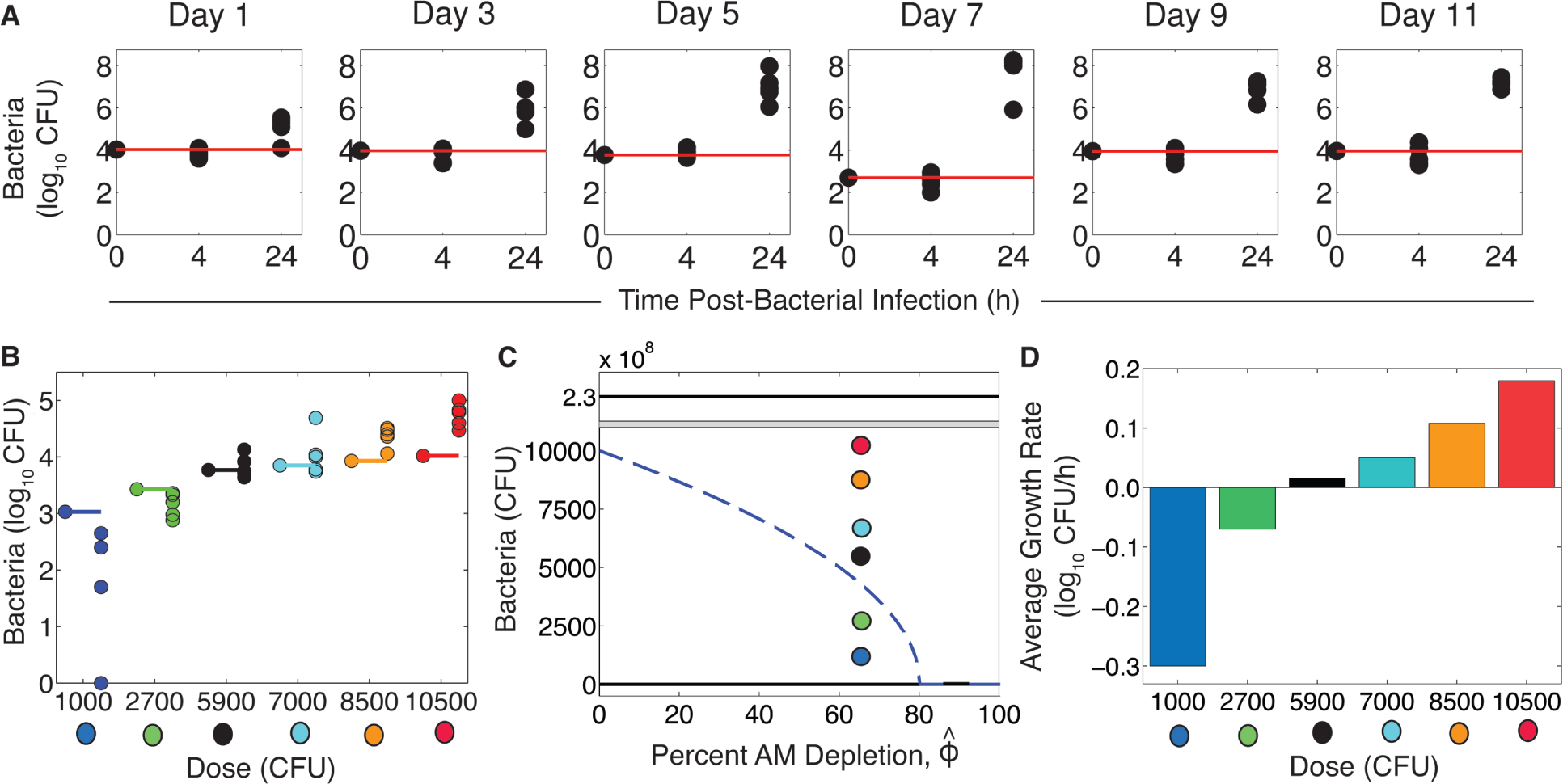
Bacterial growth for doses above the threshold. (A) Bacterial loads (log10 CFU) at 4h and 24h pbi for PR8 infection followed by D39 at the indicated coinfection time (1, 3, 5, 7, 9, or 11d pii) at doses above the threshold (Table 2). (B) Bacterial loads (log10 CFU) at 4h pbi for PR8 infection followed by D39 at 5d pii at the indicated dose (single dots on the left) with (C) the corresponding location relative to the threshold and (D) the average log10 bacterial growth rate from 0h to 4h. Colored lines indicate the inoculum size and each dot represents an individual mouse (5 mice/group).

To examine bacterial growth when AMs are depleted by a source other than virus, we gave naive mice clodronate-liposomes, which effectively reduce the AM population in the lung through cell death^27^. At 4h post-clodronate (pc), the AM population is reduced by ~75-80% (Figure 5A). Because this value is close to the critical level of AMs (Equation (1)), we infected groups of mice at 4h pc with either 1 CFU, 10 CFU, or 100 CFU. At 1 CFU, we detected bacteria in 1/5 mice at 4h pbi. At 10 CFU and 100 CFU, bacterial loads were higher than the inoculum in all 5 mice (p*>*0.05 and p<0.005, respectively) (Figure 5B).

**Figure 5.**
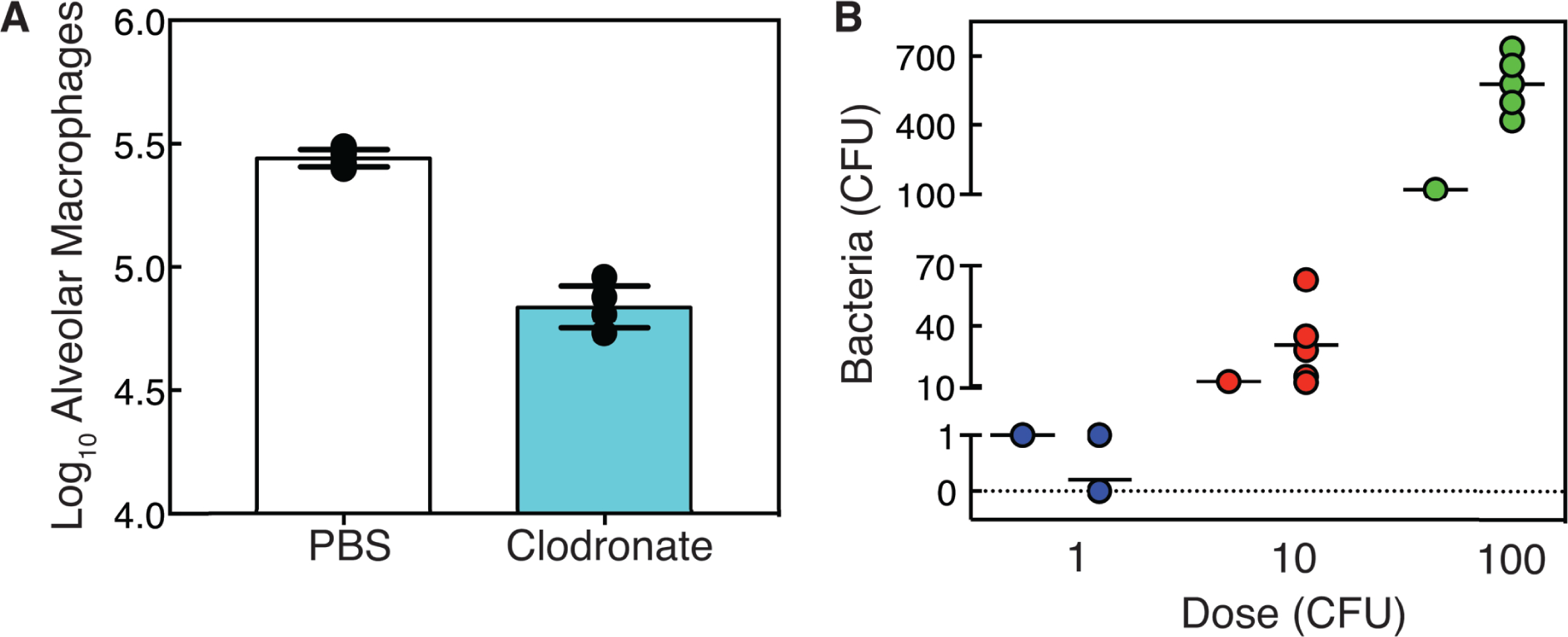
Bacterial growth after depletion of AMs by clodronate-liposome induced cell death. (A) Absolute number of AMs at 4h for mice given either PBS (white) or clodronate-liposomes (cyan). Bars are the average values, lines are the standard deviation, and each dot represents an individual mouse (5 mice/group). (B) Bacterial growth (CFU) at 4h pbi for mice given clodronate-liposomes and infected with D39 at 4h pc at the indicated dose. Black lines are the average value, single dots on the left indicate the inoculum size, and each dot represents an individual mouse (5 mice/group).

## Discussion

Secondary bacterial infections increase the severity of influenza-associated illnesses and the mortality rates during influenza pandemics^3-6^. However, not every IAV infection results in a bacterial invasion and only a proportion of successful coinfections lead to severe pneumonia^23, 24^. The probability of pneumonia manifesting during influenza is multifactorial and may involve several pathogen and host characteristics, including viral and bacterial strain, dose, and host immune status (reviewed in^7-13^). Identifying which factors promote coinfection, how each affects the kinetics, the pathogenicity, and the likelihood of bacterial pneumonia following influenza, and how they are interrelated could identify new targets for treating or preventing secondary infections.

Our analysis and experiments pinpoint the way in which bacterial dynamics vary depending on the bacterial dose and the AM population size, and identified the combinations of these two factors that lead to distinct phenotypes. In particular, we showed that an early clearance phenotype exists for doses below an initial dose threshold, a growth phenotype for doses sufficiently above the threshold, and a dichotomous phenotype for doses close to the threshold (Figures 3-5). We also show that the relationship between bacterial dose and the level of AM depletion is independent of what causes the reduction in AMs (e.g., virus infection via an unknown mechanism (Figures 3-4) or clodronate-liposomes via cell death (Figure 5)) and that 1-10 CFU is sufficient to yield bacterial growth when the depletion is ~80% (Figure 5B). The nonlinearity of the relationship between dose and AM depletion is important and non-intuitive, even with our previous knowledge that these two effects are related^16, 20-22^ and that they both influence coinfection kinetics^16-19^. Knowing how the combination of two factors work together and quantifying the conditions that yield each of the outcomes in addition to the trajectory of bacterial titers is crucial to understanding coinfection kinetics. It also allows us to predict the conditions given the behavior and vice versa.

For example, we observed that initial bacterial clearance yields larger heterogeneity in bacterial loads as the infection progresses (Figures 3A, 4A). This knowledge has given us insight into the dynamics observed in other datasets that were previously unexplained. We noted two datasets in our earlier work^17^ where bacterial titers split into two groups such that ~50% were at high levels and ~50% were at low levels (Figure S3). In the first dataset (Figure S3A), groups of mice were infected with the same virus (PR8) but with different bacterial doses (10^2^ or 10^3^ CFU) 7d pii^17^. For the larger dose (10^3^ CFU), little heterogeneity was observed in the data and bacterial growth at 4h pbi was at or above the inoculum (Figure S2B). This is consistent with a dose that is above the threshold. Comparatively, the lower dose (10^2^ CFU) resulted in some mice with high titers and some mice with low titers, which indicates a dose that is close to or below the threshold (Figure S3A). In the second dataset (Figure S3B), mice were infected with different viruses (PR8 or PR8-PB1-F2(1918)) and the same bacterial dose (10^3^ CFU) 7d pii^17^. Bacterial titers for a coinfection with the PR8-PB1-F2(1918) virus split into two groups (Figure S3B), which is consistent with an AM:dose ratio closer to the threshold, whereas little heterogeneity was observed for coinfection with the PR8 virus (Figure S2B). We hypothesize that this is due to a lower level of AM depletion caused by the PR8-PB1-F2(1918), which may be connected to the lower viral titer at 7d pii when the bacterial infection was initiated (Figure S3B inset). This is also in accordance with our model prediction that differences in bacterial dynamics were due to the antecedent viral infection^17^.

With only a short window where the surviving AMs are able to control bacterial growth, the opportunity for successful treatment while the immune system has the upper hand may be limited. Our results indicate that one preventative strategy (replenishment of AMs) and one therapeutic strategy (early reduction in bacterial loads), used separately or in combination, could be effective because the ratio of AMs to bacteria is the critical quantity that needs to be increased to abrogate secondary pneumococcal pneumonia. Reducing the transmitted dose is ideal, but this is difficult to control in practice. To prevent a coinfection, the AM population can be partially restored via immune modulators such as granuloctye macrophage colony stimulating factor (GM-CSF)^18^. In a study where mice were treated with recombinant GM-CSF (rGM-CSF) at ‐1d and 1d pii, the AM population increased by ~20% 2d after treatment (3d pii) and bacterial clearance in the first 3h after inoculation at this time point improved by ~14%^18^. Further, pneumonia was reduced from 100% to 40% and 2 out of 10 of the treated mice were able to resolve the infection within 3h compared to 0 in the untreated group. These results are consistent with those presented here, where a decrease in dose and thus an increase in the distance from the threshold results in faster clearance and resolution in some mice. Thus, similar outcomes will manifest through therapeutically decreasing AM depletion or decreasing bacterial loads, however the therapeutic requirement may change because of the nonlinearity of the relationship. A more effective therapeutic approach may be to use antibiotics, which limit bacterial replication, either prophylactically or as an early treatment together with rGM-CSF.

The importance of the AM:bacteria ratio suggests that the bacterial dose should be chosen carefully when designing pneumococcal or influenza-pneumococcal infection experiments so that misinterpretation of the results is avoided. For example, if a particular mouse strain has a greater number of AMs at baseline, a larger dose would be needed to examine a pneumococcal infection because a lower dose would be immediately cleared and the animal could be falsely regarded as protected. Further, sampling time should also be selected cautiously as important dynamics early in infection may be missed. However, based on our results, comparing the bacterial inoculum size to the final size and/or examining the heterogeneity in titers as we did here will help clarify where the experimental conditions are located along the AM:bacteria plane (Figure 2A).

Until recently, it was unknown why the morbidity and mortality from influenza-pneumococcal coinfection is maximal at 7d pii^19^. Connecting the strength of bacterial growth to the depletion of AMs, which is maximal at 7d pii, provides the key to understanding coinfection kinetics^17, 18^. Our work here quantitatively couples bacterial dose to these components and demonstrates that at least two simultaneous events (ample depletion combined with sufficient dose) are necessary for a coinfection to establish. Because both AM depletion and the dose effect are dynamic and likely dependent on other factors (e.g., AM depletion changes with viral strain), the probability of both events coinciding may be low. While a different virus strain is unlikely to impact the relationship we established between bacterial dose and AM depletion, particularly given that we found the relationship is consistent for other mechanisms of AM depletion (e.g., cell death via clodronate-liposomes (Figure 5)), strain dependent AM depletion may help to explain why only a proportion of influenza-infected individuals experience complications from secondary bacterial infections and why coinfections are less prevalent during seasonal epidemics^28^, where infections tend to be less severe compared to pandemic strains and may result in less AM depletion. When seasonal strains are in circulation, transmission of highly concentrated small particles (<10um), which are thought to deposit bacteria more readily into the lower airways (reviewed in^29^), may be required. In contrast, less concentrated and/or larger particles may be sufficient to establish a bacterial infection with more virulent strains of IAV.

Extending our results to predict the time scale at which the pneumonia progresses, the intensity of pneumonia, and the probability of survival may be more complicated. Our mathematical model (Equations (Equations (2)-(6)) is able to predict bacterial titer kinetics for a coinfection 7d pii and for infectious doses above or close to the initial dose threshold in which rapid growth ensues immediately^17^. We interpret the close fit of the model to the data (Figure S2) to mean that additional clearance mechanisms (e.g., neutrophils), which are currently excluded from the model, are ineffective. In the case where bacterial loads reach maximum levels (*>*10^8^ CFU), mortality occurs in 100% of mice within 48-72h pbi. The exact timing is dose-dependent and seems to be related to the rate at which this upper limit is achieved. However, predicting severity, outcome, and bacterial titers for infections with reduced doses, where bacterial titers may increase but remain low, is more challenging. We previously hypothesized that neutrophils may have a larger role in this context^17^. Understanding the inverse relationship between the rates of early clearance (0→4h) and later growth (4→24h) (Figs. 3B-C) does aid our ability to predict the resulting bacterial burden, but more complex and time-dependent dynamics, including inflammation and tissue damage, likely contribute to pathogenicity without significantly impacting bacterial loads.

Indeed, several other immune responses are elevated, dysfunctional, or otherwise altered during bacterial coinfection after influenza (reviewed in^7-13^). For example, subsequent clearance mechanisms (>4h pbi), such as neutrophils and macrophages, undergo influenza-induced apoptosis, become dysfunctional, and have reduced chemotactic and phagocytic functions^15, 30-34^. It is unclear the extent to which the inflated cytokine response (e.g., type I and II interferons (IFN-*α*,*β*,*γ*), TNF-*α*, IL-1*β*, IL-6, IL-12, and IL-10^16, 32, 35-48^) facilitates these changes. While our coinfection model does not explicitly account for these cytokines, the model’s ability to accurately estimate AM loss with *ϕ̂*(*V*) indicates a limited role for cytokine-mediated functional inhibition of AMs. However, the cytokine response may be related to the observed viral rebound (Figure S2A), which our model suggests is independent of AM depletion and due to a bacterial induced increase in virus production/release (*â*;(*P*))^17^. The increase in virus may be facilitated by the inhibition of IFN^49^ resulting from bacterial adherence to virus-infected cells^15, 49, 50^. With the resulting heightened state of inflammation and severe lung damage, determining how each of these events is interrelated, dose-dependent, and connected to AM depletion is critical to finding new therapeutic approaches.

Analyzing coinfection kinetics with a mathematical model provides a means to quantify and simultaneously assess multiple infection characteristics. This method allows us to make meaningful predictions about the processes altered by each pathogen, even when exact mechanisms are unknown. It also permits *in silico* experiments for systems where data is difficult to obtain (e.g., in humans), and aids experimental design to test specific predictions. Carrying out targeted experimental studies based on analytical results, as done previously^18, 49^ and here, provides new biological insight about the underlying mechanisms and pinpoints improvements that can be made to our analysis. It is only with these improvements that we will be able to further examine the interactions between influenza, pneumococcus, and the host with new models that assess other immune components. Determining the circumstances that lead to severe bacterial infections during influenza and quantifying in detail how epidemiological factors (e.g., transmission dose) and host immune status (e.g., AM depletion) work together provides important clinical insight into the threat these pathogens pose to public health. Further establishing how other pathogen (e.g., strain, viral dose) and host (e.g., neutrophils, cytokines) factors are related and contribute to other infection characteristics (e.g., probability of pneumonia and disease progression/severity) will aid the development of therapies that prevent or treat these diseases.

## Methods

### Use of Experimental Animals

All experimental procedures were approved by the Animal Care and Use Committee at SJCRH under relevant institutional and American Veterinary Medical Association guidelines and were performed in a Biosafety level 2 facility that is accredited by AALAAS.

### Influenza-Pneumococcal Coinfection Model

We previously developed a model to describe influenza-pneumococcal coinfection kinetics^17^. Briefly, the model couples single infection models for influenza virus^51^ and pneumococcus^20^ and includes terms that describe their interactions^17^. Five populations are tracked: susceptible epithelial (“target”) cells (*T*), two classes of infected cells (*I*1 and *I*_2_), virus (*V*), and bacteria (*P*).

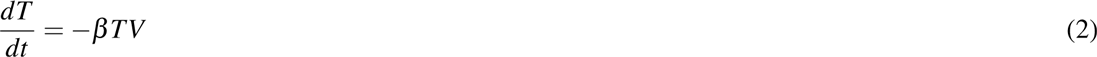

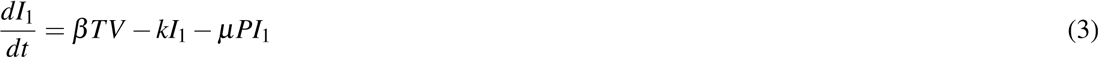

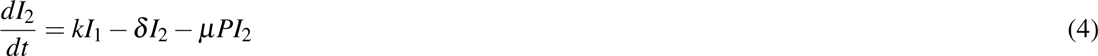

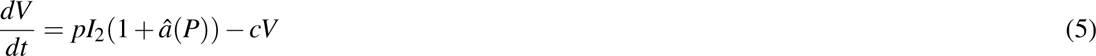

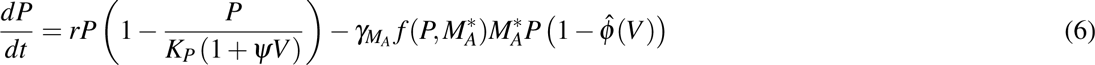

Target cells become infected with virus at rate *βV* per cell. Once infected, these cells enter an eclipse phase (*I*1) at rate *k* per cell before transitioning to produce virus at rate *p* per cell (*I*_2_). Virus is cleared at rate *c* and virus-producing infected cells (*I*_2_) are cleared at rate *δ*. Bacteria replicate logistically with maximum rate *r* and tissue carrying capacity of *K_P_*. Alveolar macrophages (*M_A_*) phagocytose bacteria at rate *γM_A_f*(*P, M_A_*) per cell. This rate decreases as the number of pneumococci increase according to *ϕ* (*P, M_A_*) = *n*^2^*M_A_*/(*P*^2^ + *n*^2^*M_A_*), where each AM is only able to phagocytose a maximum of *n* bacteria. Virus further decreases this clearance rate according to *ϕ̂*(*V*) = *ϕV*/(*K_PV_* + *V*). This term was shown to be the driving mechanism facilitating bacterial invasion^17^ and matches the percentage of AM depletion^18^. Once bacteria invade, virus production/release from infected epithelial cells (*pI*2) is increased by a factor of *â*(*P*) = *aP^z^*. This term was shown to be the driving mechanism resulting in a viral rebound^17^ and may result from IFN inhibition as a consequence of bacterial attachment to infected cells^17, 49^. The model also assumes that virus infection increases the tissue carrying capacity (*ψV*), which may facilitate bacterial adhesion to cells, and that bacteria increase infected cell death (*μP*). However, these two effects were shown to have minimal influence on the dynamics^17^. Altering other processes in the model, such as the rates of viral infection (*βV*) and clearance (*c*), produced minimal effects on model dynamics. The model schematic is shown in Figure S1, the model fits to lung viral and bacterial titers from groups of mice infected 7d after influenza A/Puerto Rico/8/34 (H1N1) (PR8) with PBS or pneumococcal strain D39 are shown in Figure S2, and the model parameters are in Table S1^17^.

### Derivation of the Initial Dose Threshold

We used standard steady state and bifurcation analyses on the coinfection model to derive the initial dose threshold. Setting Equations (2)-(6) equal to zero and solving yields 4 equilibria: a disease-free state (0,0,0,0,0) and three states with non-zero bacterial levels (0,0,0,0,*P*^*^). The non-zero *P*^*^ values satisfy *P*^3^ + *BP*^2^ + *CP* + *D* = 0 and are defined by

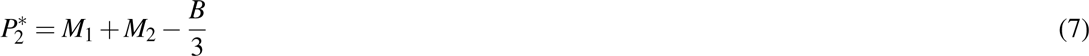

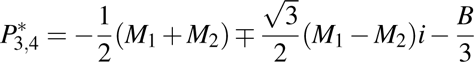

where

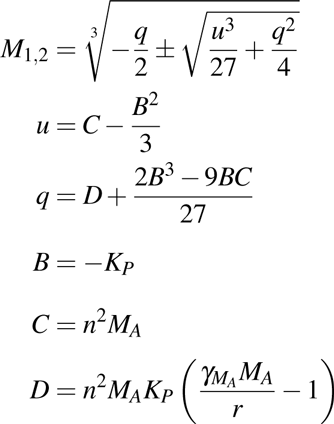

For the parameter values in Table S1, three real solutions exist because 
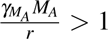
 and the discriminant, Δ = *B*^2^*C*^2^ − 4*B*^3^*D* − 4*C*^3^ + 18*BCD* − 27*D*^2^, is positive. One of the equilibria 
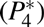
 is excluded because it is negative and not biologically relevant. Both 
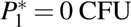
 and 
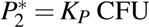
 are stable while 
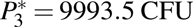
 is unstable. The unstable state 
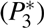
 indicates a threshold such that bacterial growth reaches the maximum carrying capacity when 
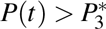
, while clearance occurs for 
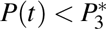
. Because *V*^*^ = 0, these equilibria are equivalent to those in the single infection models and 
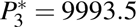
 corresponds to the initial dose threshold in the absence of an antecedent viral infection^20^.

However, virus is non-zero (*V*(*t*) > 0) at the initiation of the bacterial infection and the model solution, which is influenced by changes in the coinfection parameters^17^, is then perturbed away from its steady state. Re-solving for the equilibria yields non-zero state defined by Equation (7) but with

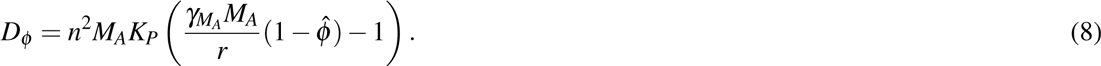

The two stable states remain unchanged (
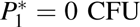
 and 
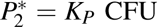
). The unstable, positive root, 
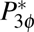
, is dependent on *ϕ̂* and real if 
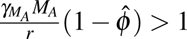
 (i.e., *D_ϕ_* > 0). However, the root turns complex when 
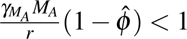
 (i.e., *D_ϕ_* < 0). The point where 
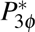
 switches from being a real root to a complex root with real part less than 1 is found by solving *D_ϕ_* = 0 for *ϕ̂*, which gives the critical value in Equation (1). The equilibria are plotted against *ϕ̂* in Figure 2A.

### Mice

Adult (6 week old) female BALB/cJ mice were obtained from Jackson Laboratories (Bar Harbor, ME). Mice were housed in groups of 5 mice in high-temperature 31.2cm × 23.5cm × 15.2cm polycarbonate cages with isolator lids. Rooms used for housing mice were maintained on a 12:12-hour light:dark cycle at 22 ± 2°C with 50% humidity in the biosafety level 2 facility at St. Jude Children’s Research Hospital (Memphis, TN). Prior to inclusion in the experiments, mice were allowed at least 7 days to acclimate to the animal facility such that they were 7 weeks old at the time of infection. Laboratory Autoclavable Rodent Diet (PMI Nutrition International, St. Louis, MO) and autoclaved water were available ad libitum. All experiments were performed under an approved protocol and in accordance with the guidelines set forth by the Animal Care and Use Committee at St. Jude Children’s Research Hospital.

### Infectious Agents

All experiments were done using the mouse adapted influenza A/Puerto Rico/8/34 (H1N1) (PR8) and type 2 pneumococcal strain D39 that was transformed with the lux operon (Xenogen) to make it bioluminescent^52^.

### Macrophage Depletion by Clodronate-Liposomes

Clodronate-liposomes (5mg/ml, clodronateliposomes.org) were delivered intranasally in 100ul.

### Infection Experiments

The viral infectious dose (TCID50) was determined by interpolation using the method of Reed and Muench^53^ using serial dilutions of virus on Madin-Darby canine kidney (MDCK) cells. The bacterial infectious dose (CFU) were counted for serial dilutions of bacteria on tryptic soy-agar plates supplemented with 3% (vol/vol) sheep erythrocytes. Inocula were diluted in sterile PBS and administered intranasally to groups of 5 mice lightly anesthetized with 2.5% inhaled isoflurane (Baxter, Deerfield, IL) in a total volume of 100ul (50ul per nostril). Mice were inoculated with 50 TCID50 PR8 at day 0 and with D39 at 1, 3, 5, 7, 9, or 11d pii at the doses listed in Table 2. Clodronate-liposome treated mice were inoculated with either PBS or D39 at 4h post-treatment at a dose of 1, 10, or 100 CFU in 100ul. Mice were weighed at the onset of infection and each subsequent day for illness and mortality. Mice were euthanized if they became moribund or lost 30% of their starting body weight. We repeated each experiment at least one time to ensure reproducibility.

### Lung Titers

Mice were euthanized by CO2 asphyxiation. Lungs were aseptically harvested, washed three times in PBS, and placed in 500ul PBS. Lungs were mechanically homogenized using the Ultra-Turrax T8 homogenizer (IKA-werke, Staufen, Germany). Lung homogenates were pelleted at 10,000 rpm for 5 minutes and the supernatants were used to determine the bacterial titer for each set of lungs using serial dilutions on tryptic soy-agar plates supplemented with 3% (vol/vol) sheep erythrocytes.

### Flow Cytometric Analysis of Alveolar Macrophages

After euthanasia by CO2 inhalation, whole lungs were harvested, digested with collagenase (1mg/ml, Sigma C0130), and physically homogenized by syringe plunger against a 40um cell strainer. Cell suspensions were centrifuged at 4°C, 500xg for 7 min. Following red blood cell lysis, cells were washed in MACS buffer (PBS, 0.1M EDTA, 0.01M HEPES, 5mM EDTA and 5% heat-inactivated FBS) and counted with trypan blue exclusion using a Cell Countess System (Invitrogen, Grand Island, NY). Flow cytometry (LSRII Fortessa; Becton Dickinson, San Jose, CA) was performed on the cell pellets after incubation with 200ul of 1:2 dilution of Fc block (human-*γ* globulin) on ice for 30 min, followed by surface marker staining with anti-mouse antibodies: CD11c (eFluor450, eBioscience), CD11b (Alexa700, BD Biosciences), Ly6G (PerCp-Cy5.5, Biolegend), Ly6C (APC, eBioscience), F4/80 (PE, eBioscience), CD3e (PE-Cy7), CD4 (PE-Cy5, BD Biosciences), CD8a (BV605, BD Biosciences), DX5 (APC-Cy7, Biolegend) and MHC-II (FITC, eBioscience). The data were analyzed using FlowJo 10.0.8 (Tree Star, Ashland, OR) where viable cells were gated from a forward scatter/side scatter plot and singlet inclusion (see Figure S4). Following neutrophil exclusion (Ly6Ghi), AMs were gated as CD11chiF4/80hiCD11b^−^. The absolute numbers of different cell types were calculated based on viable events analyzed by flow cytometry as related to the total number of viable cells per sample. We obtain the range of percentage of AMs depleted by normalizing the absolute number of AMs to the absolute number of AMs in a naive mouse (Figure 5).

### Alveolar Macrophage Data

The AM data used in this study is from Ref.^18^ (Copyright 2013. The American Association of Immunologists, Inc.). In brief, groups of five 6-8 week old female BALB/cJ mice were infected intranasally with the PR8 virus. Bronchoalveolar lavage fluid (BALF) was collected and whole lungs post-lavage were harvested. Cells were analyzed by flow cytometry and AMs were gated as Ly6G-F4/80^hi^CD11c^hi^CD11b^−^. Here, we use the AM data from the lung and obtain the percentage of AMs depleted by normalizing the absolute number of AMs at each time point to the average number of AMs in a naive mouse (Figure 2B).

### Statistical Comparisons

We used the statistical programming language R^54^.T-tests were used to determine significance of differences of bacterial titers. A p < 0.05 was considered significant for these comparisons.

## Author Contributions

A.M.S conceived the concept, planned the experiments, and wrote the manuscript. A.M.S. and A.P.S. performed the experiments. A.M.S. analyzed the model and data.

## Acknowledgments

This work was supported by NIH grants AI100946 and AI125324 and by ALSAC. We thank Fred Adler, Jon McCullers, and Alan Perelson for their helpful comments.

